# Identification of *Photorhabdus* symbionts by MALDI-TOF mass spectrometry

**DOI:** 10.1101/2020.01.10.901900

**Authors:** Virginia Hill, Peter Kuhnert, Matthias Erb, Ricardo A. R. Machado

## Abstract

Species of the bacterial genus *Photorhabus* live in a symbiotic relationship with *Heterorhabditis* entomopathogenic nematodes. Besides their use as biological control agents against agricultural pests, some *Photorhabdus* species are also a source of natural products and are of medical interest due to their ability to cause tissue infections and subcutaneous lesions in humans. Given the diversity of *Photorhabdus* species, rapid and reliable methods to resolve this genus to the species level are needed. In this study, we evaluated the potential of matrix-assisted laser desorption/ionization time-of-flight mass spectrometry (MALDI-TOF MS) for the identification of *Photorhabdus* species. To this end, we established a collection of 54 isolates consisting of type strains and multiple field strains that belong to each of the validly described species and subspecies of this genus. Reference spectra for the strains were generated and used to complement a currently available database. The extended reference database was then used for identification based on the direct transfer sample preparation method and protein fingerprint of single colonies. High discrimination of distantly related species was observed. However, lower discrimination was observed with some of the most closely related species and subspecies. Our results, therefore, suggest that MALDI-TOF MS can be used to correctly identify *Photorhabdus* strains at the genus and species level, but has limited resolution power for closely related species and subspecies. Our study demonstrates the suitability and limitations of MALDI-TOF-based identification methods for the assessment of the taxonomical position and identification of *Photorhabdus* isolates.

**Impact Statement:** Species of the bacterial genus *Photorhabus* live in close association with soil-born entomopathogenic nematodes. Under natural conditions, these bacteria are often observed infecting soil-associated arthropods, but under certain circumstances, can also infect humans. They produce a large variety of natural products including antibiotics, insecticides, and polyketide pigments that have substantial agricultural, biotechnological and medical potential. In this study, we implement MALDI-TOF MS-based identification method to resolve the taxonomic identity of this bacterial genus, providing thereby a rapid identification tool to understanding its taxonomic diversity to boost scientific progress in medical, agricultural, and biotechnological settings.

## Introduction

Species of the genus *Photorhabdus* live in a close symbiotic association with *Heterorhabditis* entomopathogenic nematodes (1). Given their biosynthetic capacity and ability to produce a large array of specialized metabolites and proteins, and their ability to infect humans and arthropods, *Photorhabdus* species are of biotechnological, medical, and agricultural interest (2–5). Understanding their taxonomic diversity is an important step towards minimizing human health risks and maximizing the agricultural and biotechnological potential of *Photorhabdus* species.

In natural ecosystems, *Photorhabdus* species are carried by entomopathogenic nematodes in their intestines. Entomopathogenic nematodes colonize soil-born arthropods, and release these bacteria in the hemocoel of their prey (6, 7). Bacteria reproduce, produce toxins, immune suppressors, and lytic enzymes, cause septicemia, toxemia, and in many cases kill the infected organism (8). Consequently, these organisms are broadly used as biological control agents to combat agricultural pests (9, 10, 4, 11, 12). Under some particular cases, however, certain *Photorhabdus* species as *Photorhabdus asymbiotica* have been reported to infect humans and cause local tissue infections and subcutaneous nodules (13–16, 3).

Possibly due to their particular lifestyle, *Photorhabdus* species produce an arsenal of secondary metabolites (17–19, 2, 20–22). These metabolites act as virulence factors to kill their prey, symbiosis factors to support the growth of their nematode host, and/or antimicrobial compounds that limit the proliferation of microbial competitors (23–27, 5, 28, 20). Apart from their ecological importance, these metabolites are also valuable for biotechnology. For instance, 3,5-dihydroxy-4-isopropylstilbene, produced by *Photorhabdus* sp. C9, shows antifungal activities against important medical and agricultural fungi like *Aspergillus flavus* and *Candida tropicalis* (29). Another example is carbapenem, an important broad-spectrum β-lactam antibiotic produced by *Photorhabdus luminescence* strain TT01 (30).

Due to their importance as biocontrol agents, human pathogens, and bio-factories, substantial efforts have been made to understand the diversity of the *Photorhabdus* bacterial group. For this, several collection campaigns have been set around the world which have yield many different isolates (31–44). In addition, several methods for the identification of these isolates have been developed and implemented. In medical cases for instance, bacteria isolated from diseased human tissues were identified using classical microbiological methods such as characterization of colony morphology and VITEK 2 Gram-negative identification card-based biochemical tests (45, 15). Unfortunately, these methods misleadingly assigned the causing agent to other bacterial species (46). Routine automated mass spectrometry methods failed to identify the potential disease-causing agent, because *Photorhabdus* spectra were absent from databases. Finally, 16S rRNA gene sequencing had to be performed to properly identify the bacterium that caused the cutaneous lesions (47). Other methods such as restriction fragment length polymorphism-PCR (RFLP-PCR) were used, but also proved to be of very limited taxonomic value (48, 49). Later, multi locus sequence analysis (MLSA) was found to be a useful tool for the taxonomic description of *Photorhabdus* species (50). However, with an increasing number of available strains and due to the high taxonomic complexity of this bacterial group, whole-genome based methods were shown to be particularly suitable to resolve the phylogenetic relationship of especially closely related *Photorhabdus* species and subspecies (34, 33). Nonetheless, these methods are laborious and do not allow for a rapid identification. The above mentioned limitations might be overcome by MALDI-TOF-based identification techniques (51–55).

In this study, we evaluate the possibility to use MALDI-TOF to rapidly resolve the taxonomic identity of *Photorhabdus* species. To this end, we established an experimental collection of *Photorhabdus* isolates including type and several field strains of all the validly described species of this genus. We then created main spectra libraries and constructed MALDI-TOF MS-based dendrograms. The results of our study highlight the possibilities and limitations of MALDI-TOF MS for the identification of *Photorhabdus* species.

## Materials and methods

### Bacterial strains

The 54 bacterial strains included in this study were either part of our *in-house* collection, were kindly provided by different collaborators, or were acquired from biological resource centers (Czech Collection of Microorganisms, CCM, or the Leibniz Institute DSMZ-German Collection of Microorganisms and Cell Cultures, DSMZ). All the bacterial species have been previously isolated from their nematode host or from soft wounds of human patients as described (56, 33, 34, 50, 38–44, 1, 37).

### Generation of phylogenetic trees

To identify the *Photorhabdus* bacterial strains as a baseline for the MALDI-TOF experiment, we followed the procedure described by Machado *et. al*. (33, 34). Briefly, bacterial genomic DNA was extracted using the GenElute Bacterial Genomic DNA Kit (Sigma-Aldrich) following the manufacturer’s instructions. The 16S rRNA gene was amplified by PCR using the following primers: 8F AGAGTTTGATCCTGGCTCAG and 1492R CGGTTACCTTGTTACGACTT. PCR products were separated by electrophoresis in a 1 % TAE-agarose gel stained with GelRed nucleic acid gel stain (Biotium), gel-purified (QIAquick gel purification Kit, Qiagen) and sequenced by Sanger sequencing (Microsynth). Obtained sequences were manually curated, trimmed and used to reconstruct evolutionary histories using the Neighbor-Joining method (57). The optimal tree with the sum of branch length = 0.20174092 is shown. The percentage of replicate trees in which the associated taxa clustered together in the bootstrap test (100 replicates) are shown next to the branches (58). The tree is drawn to scale, with branch lengths in the same units as those of the evolutionary distances used to infer the phylogenetic tree. The evolutionary distances were computed using the Kimura 2-parameter method (59) and are in the units of the number of base substitutions per site. There were a total of 1166 positions in the final dataset. Evolutionary analyses were conducted in MEGA7 (60). Graphical representation and edition of the phylogenetic tree were performed with Interactive Tree of Life (version 3.5.1) (61, 62). Whole-genome-based phylogenetic tree was adapted from (34).

### Generation of main spectra

Main spectra (MSP) were generated on a microflex™LT (Bruker Daltonik GmbH, Bremen, Germany) as described (63). For this, bacteria were grown from glycerol stocks on Luria Bertani (LB) plates at 28°C for 28 hours. Proteins were then extracted following the standard formic acid-based method recommended by the manufacturer (Bruker Daltonik GmbH). Briefly, a few single bacterial colonies were dissolved in 300 µl of pyrogen free water by vortexing and 900 µl of absolute ethanol were added to the solution. After centrifugation (2 min, 15’000 rpm), the supernatant was discarded. After a second centrifugation step and removing the remaining ethanol, the pellet was air dried for 2-3 min. Pellets were then resuspended in 30 µl of 70% formic acid (Sigma Aldrich, Germany). Subsequently, 30 µl of acetonitrile (Fluka analytical, Germany) were added and mixed by pipetting followed by centrifugation for 2 min. at 15’000 rpm. One µl of the resulting supernatant was transferred to the MALDI target plate in eight replicates and let dry at room temperature. Then, 1 µl of matrix (α-Cyano-4-hydroxycinnamicacid, HCCA, CAS Number 28166-41-8, Sigma-Aldrich, Switzerland) was added. Each spot was then measured in triplicate to obtain 24 spectra per strain using the MBT_AutoX method of flexControl software. The generated spectra were visually inspected and edited in the flexAnalysis software according to Bruker recommendations. Individual spectra diverging from the cohort core, i.e. differing by more than 500 ppm, were deleted. A minimum of 20 spectra per strain were then used for the generation of MSP in the MBT Compass Explorer 4.1 (Bruker) using standard settings. The MSP of each strain was added to the project library used for identification. Newly generated MSP were entered to the project database and were used for identification and to generate a dendrogram using the correlation distance measure with the average linkage algorithm and a threshold value for a single organism of 300 in MBT Compass Explorer 4.1. The MBT Compass Library 4.1 currently contains 3000 bacterial species of 540 genera.

### Diagnostic identification

To validate the suitability of our newly established main spectra database for the identification of *Photorhabdus* species, several *Photorhabdus* strains with known taxonomic identities were tested. To this end, the bacterial strains were grown for 28 hours at 28°C on LB media. Single colonies were picked with toothpicks, transferred onto the MALDI-TOF target plate, dried at room temperature and mounted with 1 µl of HCCA matrix. Identification of *Photorhabdus* strains was performed by comparing the resulting spectra against the extended Bruker database, including the newly generated *Photorhabdus* MSP.

## Results and discussion

A collection of 54 *Photorhabdus* strains belonging to all the 22 validly described species and subspecies was used to evaluate the suitability of MALDI-TOF to identify *Photorhabdus* species. Based on the main spectra produced from the strains, a dendrogram was generated (Figure 1). Two main clusters were observed. All strains of a species clustered together. In some cases, however, clusters composed of strains that belong to two or more species were observed. In particular, all the strains of *P. khanii* subsp. *khanii, P. khanii subsp. guanajuatensis, P. stackebrandtii, P. tasmaniensis, P. thracensis*, and *P. thracensis* form a cluster; all strains of *P. cinerea* and *P. heterorhabditis* form a cluster; all strains of *P. luminescens* subsp. *mexicana* and *P. luminescens subsp. luminescens* form a cluster; and all the strains of *P. kayaii, P. kleinii and P. bodei* formed a cluster. The MSP-based dendrogram topology barely resembled the topology of the 16S rRNA gene based phylogenetic tree (Figure 2), but closely mirrored the whole genome-based tree (Figure 3).

**Figure 1.**
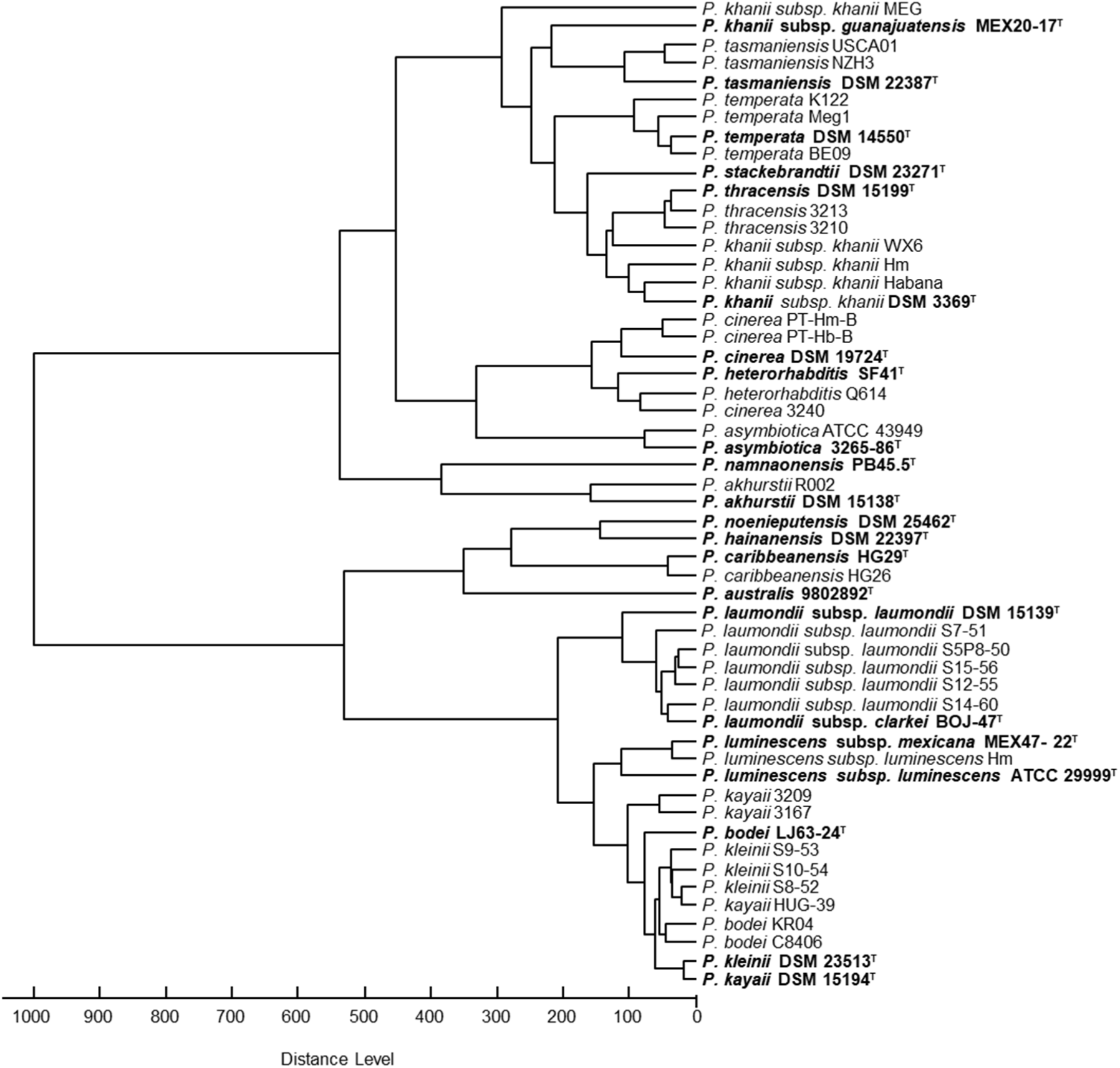
Main spectra (MSP)-based dendrogram of *Photorhabdus* strains. Type strains are indicated in bold. The distance level is normalized to a maximum value of 1000.

**Figure 2.**
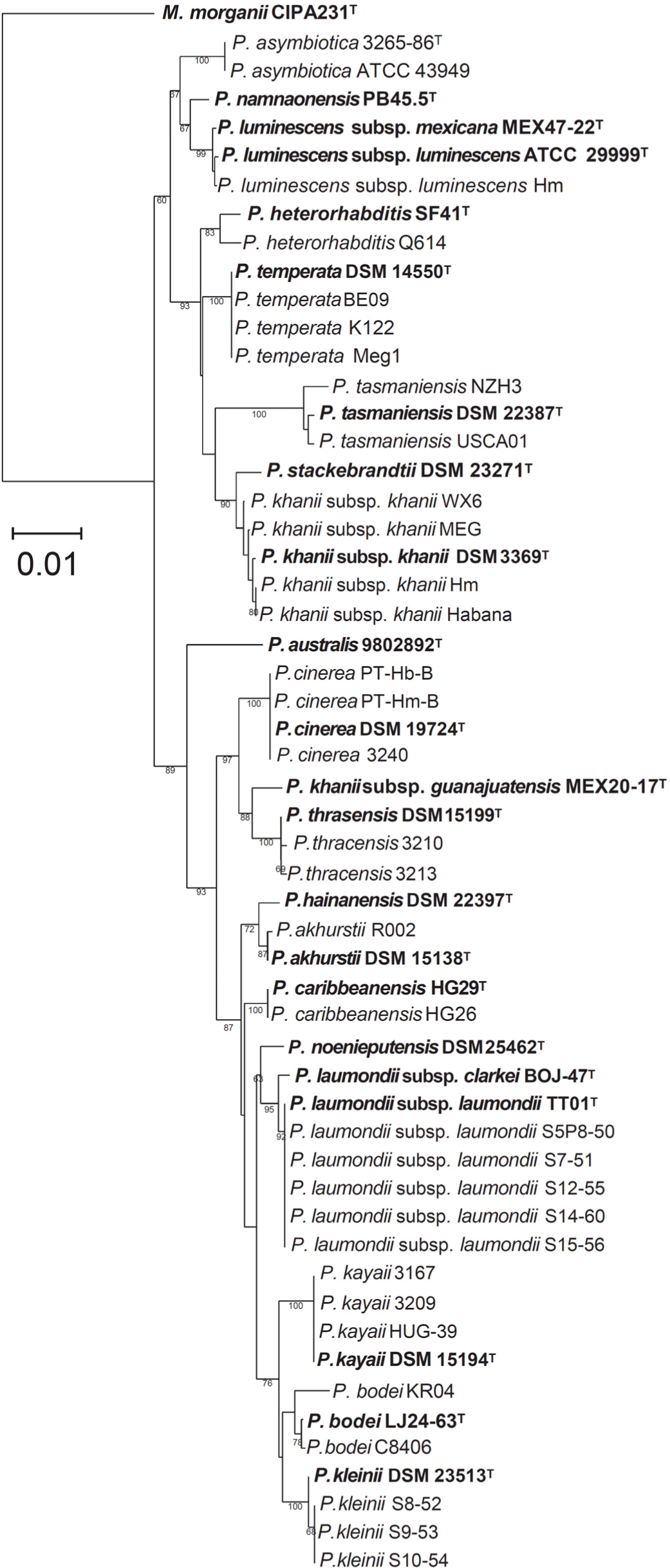
Neighbor-Joining based phylogenetic tree of *Photorhabdus* bacterial strains reconstructed from 1166 nucleotide positions of 16S ribosomal RNA gene sequences. Numbers at nodes represent bootstrap values based on 100 replications. Bar, 0.01 nucleotide substitutions per sequence position. Sequences used were deposited into the National Center for Biotechnology Information (NCBI) databank. Accession numbers are listed in Table S1.

**Figure 3.**
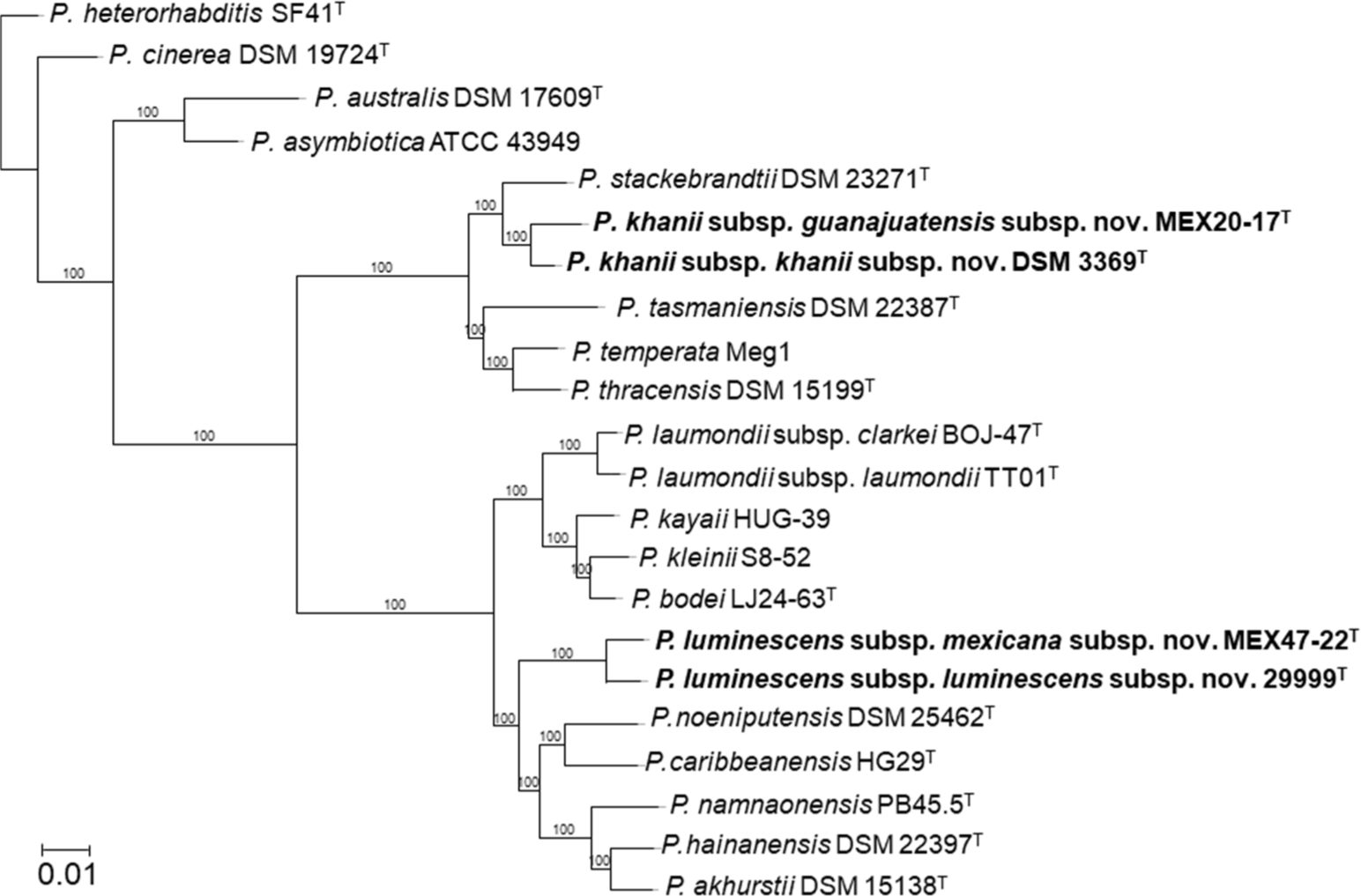
Phylogenetic reconstruction based on core genome sequences of *Photorhabdus* bacterial strains. 1662 open reading frames were analyzed. Numbers at the nodes represent SH-like branch supports. Bar, 0.01 nucleotide substitutions per sequence position.

Using identification scores, all strains could unequivocally be identified up to the genus level, with some limitations for closely related species. Twenty-five (45%) and fifteen (27%) of the analyzed strains appear either first or second, respectively, in the list of strains with best matching scores. For 96% of the strains, greater matching scores were observed with strains of their own species than with members of other species (Table 1). Only two strains *P. laumondii* subsp. *laumondii* S14-60 and *P. namnaonensis* PB45.5^T^ show similar matching scores with members of other species. We also observed that the identification scores of species that are more closely related, according to whole-genome based phylogenies, tended to be more similar than the scores of strains that are only distantly related. These effects were also observed in the MSP-based dendrogram (Table 1, Figure 1). In particular, we observed that strains of *P. laumondii subsp. laumondii, P. laumondii subsp. clarkei, P. kayaii, P. kleinii and P. bodei* were normally listed within the 10 best scores when analyzing any strain belonging to these species. Similarly occurred for strains that belong to *P. luminescens subsp. luminescens, P. luminescens* subsp. *mexicana, P. noeniputensis, P. caribbeanensis, P. namnaonensis, P. hainanensis* and *P. akhurstii*. For the identification of some strains that belong to the abovementioned species, additional tests might therefore be required. In this context, citrate utilization, indole and acetoin production, and tryptophan deaminase, gelatinase and glucose oxidase activity have been shown to be particularly useful for the discrimination of *Photorhabdus* species (34).

**Table 1.** MALDI BioTyper identification score values for different *Photorhabdus* strains. Score values higher than 1.99: secure to highly probable species identification; between 1.7 and 1.99: probable genus identification; and between 0.0 and 1.69: not reliable identification. ^T^ indicates type strain.

| Species | Strain | Best match with |  |
| --- | --- | --- | --- |
|  |  | Same species | Different species |
| *P. akhurstii* | DSM 15138<sup>T</sup> | 2.26 | 2.05 (*P. caribbeanensis*) |
|  | R002 | 2.47 | 2.18 (*P. hainanensis*) |
| *P. asymbiotica* | 3265-86<sup>T</sup> | 2.38 | 1.87 (*P. australis*) |
|  | ATCC 43949 | 2.51 | 2.06 (*P. australis*) |
| *P. australis* | 9802892<sup>T</sup> | 2.44 | 1.94 (*P. cinerea*) |
| *P. bodei* | LJ63-24<sup>T</sup> | 2.54 | 2.36 (*P. kleinii*) |
|  | KR04 | 2.38 | 2.29 (*P. kleinii*) |
|  | C8406 | 2.46 | 2.32 (*P. kleinii*) |
| *P. caribbeanensis* | HG29<sup>T</sup> | 2.62 | 2.29 (*P. hainanensis*) |
|  | HG26 | 2.69 | 2.25 (*P. hainanensis*) |
| *P. cinerea* | DSM 19724<sup>T</sup> | 2.50 | 1.80 (*P. asymbiotica*) |
|  | 3240 | 2.63 | 2.00 (*P. asymbiotica*) |
|  | PT-Hb-B | 2.50 | 1.86 (*P. heterorhabditis*) |
|  | PT-Hm-B | 2.55 | 1.98 (*P. heterorhabditis*) |
| *P. hainanensis* | DSM 22397<sup>T</sup> | 2.49 | 2.30 (*P. noenieputensis*) |
| *P. heterorhabditis* | SF41<sup>T</sup> | 2.37 | 2.14 (*P. cinerea*) |
|  | Q614 | 2.46 | 2.17 (*P. cinerea*) |
| *P. kayaii* | DSM 15194<sup>T</sup> | 2.26 | 2.19 (*P. kleinii*) |
|  | HUG-39 | 2.55 | 2.39 (*P. bodei*) |
|  | 3167 | 2.51 | 2.26 (*P. kleinii*) |
|  | 3209 | 2.47 | 2.37 (*P. kleinii*) |
| *P. khanii* subsp. *guanajuatensis* | MEX20-17<sup>T</sup> | 2.49 | 2.17 (*P. thracensis*) |
| *P. khanii* subsp. *khanii* | DSM 3369<sup>T</sup> | 2.22 | 1.85 (*P. stackebrandtii*) |
|  | WX6 | 2.44 | 2.08 (*P. tasmaniensis*) |
|  | MEG | 2.53 | 2.11 (*P. temperata*) |
|  | Habana | 2.53 | 2.07 (*P. temperata*) |
|  | Hm | 2.36 | 1.88 (*P. thracensis*) |
| *P. kleinii* | DSM 23513<sup>T</sup> | 2.50 | 2.44 (*P. bodei*) |
|  | S8-52 | 2.47 | 2.32 (*P. bodei*) |
|  | S9-53 | 2.43 | 2.33 (*P. bodei*) |
|  | S10-54 | 2.40 | 2.37 (*P. bodei*) |
| *P. laumondii* subsp. *clarkei* | BOJ-47<sup>T</sup> | 2.40 | 1.88 (*P. luminescens* subsp. *mexicana*) |
| *P. laumondii* subsp. *laumondii* | DSM 15139<sup>T</sup> | 2.42 | 2.00 (*P. kayaii*) |
|  | S12-55 | 2.20 | 1.77 (*P. kleinii*) |
|  | S5P8-50 | 2.22 | 1.70 (*P. kayaii*) |
|  | S15-56 | 2.50 | 1.84 (*P. kayaii*) |
|  | S14-60 | 2.50 | 2.51 (*P. kleinii*) |
|  | S7-51 | 2.37 | 1.86 (*P. kayaii*) |
| *P. luminescens* subsp. *luminescens* | ATCC 29999<sup>T</sup> | 2.39 | 2.18 (*P. noenieputensis*) |
|  | Hm | 2.44 | 2.22 (*P. hainaniensis*) |
| *P. luminescens* subsp. *mexicana* | MEX47-22<sup>T</sup> | 2.56 | 2.33 (*P. noenieputensis*) |
| *P. namnaonensis* | PB45.5<sup>T</sup> | 2.33 | 2.33 (*P. luminescens* subsp. *mexicana*) |
| *P. noenieputensis* | DSM 25462<sup>T</sup> | 2.53 | 2.27 (*P. hainanensis*) |
| *P. stackebrandtii* | DSM 23271<sup>T</sup> | 2.44 | 2.01 (*P. asymbiotica*) |
| *P. tasmaniensis* | DSM 22387<sup>T</sup> | 2.53 | 2.15 (*P. temperata*) |
|  | USCA01 | 2.37 | 2.17 (*P. temperata*) |
|  | NZH3 | 2.44 | 2.22 (*P. temperata*) |
| *P. temperata* | DSM 14550<sup>T</sup> | 2.47 | 2.18 (*P. thracensis*) |
|  | Meg1 | 2.42 | 2.11 (*P. tasmaniensis*) |
|  | K122 | 2.48 | 2.14 (*P. thracensis*) |
|  | BE09 | 2.63 | 2.19 (*P. tasmaniensis*) |
| *P. thracensis* | DSM 15199<sup>T</sup> | 2.33 | 2.11 (*P. temperata*) |
|  | 3210 | 2.53 | 2.28 (*P. temperata*) |
|  | 3213 | 2.49 | 2.16 (*P. temperata*) |

## Conclusion

MALDI-TOF MS was shown to be a powerful method to identify *Photorhabdus* species at the genus level and in many cases up to the species level. Some limitations were observed for closely related species and subspecies, for which additional tests might be necessary. No special sample preparation is required as the direct transfer sample preparation method is sufficient for generating good quality spectra for comparison against available spectral databases.

## Acknowledgments

We thank the following scientists and institutions for sharing bacteria or nematode strains: Helge Bode (Goethe University Frankfurt, Germany), Louis Tisa (University of New Hampshire, United Kingdom), Sylvie Pages (INRA-University of Montpellier, France), Tamas Lakatos (National Agricultural Research and Innovation Centre, Hungary), Toyoshi Yoshiga (Saga University, Japan), Waldemar Kazimierczak (University of Lublin, Poland), Ted Turlings (University of Neuchatel, Switzerland), Pamela Bruno (University of Neuchatel, Switzerland), Juan Ma (Integrated Pest Management Centre of Hebei Province, PR China), Javad Karimi (Ferdowsi University of Mashhad, Iran), and David Clarke (University College Cork, Ireland). We thank Lisa Thönen, Anja Boss, and Evangelia Vogiatzaki (University of Bern) for their assistance with bacterial colony maintenance.

## Conflicts of interest

The authors declare that there are no conflicts of interest.

## Data availability

The MSP generated in this study will be made available to interested scientist upon reasonable request to the authors.

## Funding information

This study was supported by the University of Bern (Switzerland) and received no specific grant from any funding agency.

## Supplementary Material

**Table S1.** National Center for Biotechnology Information (NCBI) accession numbers of 16S ribosomal RNA gene sequences of all *Photorhabdus* bacterial strains used in this study.

| Species | Strain | NCBI accession number |
| --- | --- | --- |
| *P. akhurstii* | DSM 15138<sup>T</sup> | MN714266 |
|  | R002 | MN714235 |
| *P. asymbiotica* | 3265-86<sup>T</sup> | MN714241 |
|  | ATCC 43949 | MN714278 |
| *P. australis* | 9802892<sup>T</sup> | MN714240 |
| *P. bodei* | LJ63-24<sup>T</sup> | MN714238 |
|  | KR04 | MN714253 |
|  | C8406 | MN714254 |
| *P. caribbeanensis* | HG29<sup>T</sup> | MN714263 |
|  | HG26 | MN714279 |
| *P. cinerea* | DSM 19724<sup>T</sup> | MN714242 |
|  | 3240 | MN714273 |
|  | PT-Hb-B | MN714231 |
|  | PT-Hm-B | MN714232 |
| *P. hainanensis* | DSM 22397<sup>T</sup> | MN714265 |
| *P. heterorhabditis* | SF41<sup>T</sup> | MN714243 |
|  | Q614 | MN714272 |
| *P. kayaii* | DSM 15194<sup>T</sup> | MN714255 |
|  | HUG-39 | MN714239 |
|  | 3167 | MN714233 |
|  | 3209 | MN714234 |
| *P. khanii* subsp. *guanajuatensis* | MEX20-17<sup>T</sup> | MN714283 |
| *P. khanii* subsp. *khanii* | DSM 3369<sup>T</sup> | MN714246 |
|  | WX6 | MN714271 |
|  | MEG | MN714269 |
|  | Habana | MN714282 |
|  | Hm | MN714236 |
| *P. kleinii* | DSM 23513<sup>T</sup> | MN714249 |
|  | S8-52 | MN714250 |
|  | S9-53 | MN714251 |
|  | S10-54 | MN714252 |
| *P. laumondii* subsp. *clarkei* | BOJ-47<sup>T</sup> | MN714237 |
| *P. laumondii* subsp. *laumondii* | DSM 15139<sup>T</sup> | MN714256 |
|  | S12-55 | MN714259 |
|  | S5P8-50 | MN714257 |
|  | S15-56 | MN714261 |
|  | S14-60 | MN714260 |
|  | S7-51 | MN714258 |
| *P. luminescens* subsp. *luminescens* | ATCC 29999<sup>T</sup> | MN714262 |
|  | Hm | MN714276 |
| *P. luminescens* subsp. *mexicana* | MEX47-22<sup>T</sup> | MN714284 |
| *P. namnaonensis* | PB45.5<sup>T</sup> | MN714267 |
| *P. noenieputensis* | DSM 25462<sup>T</sup> | MN714264 |
| *P. stackebrandtii* | DSM 23271<sup>T</sup> | MN714245 |
| *P. tasmaniensis* | DSM 22387<sup>T</sup> | MN714244 |
|  | USCA01 | MN714281 |
|  | NZH3 | MN714280 |
| *P. temperata* | DSM 14550<sup>T</sup> | MN714247 |
|  | Meg1 | MN714277 |
|  | K122 | MN714275 |
|  | BE09 | MN714274 |
| *P. thracensis* | DSM 15199<sup>T</sup> | MN714248 |
|  | 3210 | MN714268 |
|  | 3213 | MN714270 |

## Notes

#### Summary of Updates

Species of the bacterial genus Photorhabus live in a symbiotic relationship with Heterorhabditis entomopathogenic nematodes. Besides their use as biological control agents against agricultural pests, some Photorhabdus species are also a source of natural products and are of medical interest due to their ability to cause tissue infections and subcutaneous lesions in humans. Given the diversity of Photorhabdus species, rapid and reliable methods to resolve this genus to the species level are needed. In this study, we evaluated the potential of matrix-assisted laser desorption/ionization time-of-flight mass spectrometry (MALDI-TOF MS) for the identification of Photorhabdus species. To this end, we established a collection of 54 isolates consisting of type strains and multiple field strains that belong to each of the validly described species and subspecies of this genus. Reference spectra for the strains were generated and used to complement a currently available database. The extended reference database was then used for identification based on the direct transfer sample preparation method and protein fingerprint of single colonies. High discrimination of distantly related species was observed. However, lower discrimination was observed with some of the most closely related species and subspecies. Our results, therefore, suggest that MALDI-TOF MS can be used to correctly identify Photorhabdus strains at the genus and species level, but has limited resolution power for closely related species and subspecies. Our study demonstrates the suitability and limitations of MALDI-TOF-based identification methods for the assessment of the taxonomical position and identification of Photorhabdus isolates.

